# Transcriptomic analysis of fetal membranes reveals pathways involved in preterm birth

**DOI:** 10.1101/358945

**Authors:** Silvana Pereyra, Claudio Sosa, Bernardo Bertoni, Rossana Sapiro

**Author notes:** Corresponding author: (RS).

## Abstract

Preterm birth (PTB), defined as infant delivery before 37 weeks of completed gestation, results of the interaction of both genetic and environmental components and constitutes a complex multifactorial syndrome. Transcriptome analysis of PTB has proved challenging because of the multiple causes of PTB and the numerous maternal and fetal gestational tissues that must interact to facilitate parturition. A common pathway of labor and PTB may be the activation of fetal membranes. In this work, chorioamnion membranes from severe preterm and term fetus were analyzed using RNA sequencing. A total of 270 genes were differentially expressed (DE): 252 were up-regulated and 18 were down-regulated in the severe preterm compared to the term births. We found great gene expression homogeneity in the control samples, and not in severe preterm samples. In this work, we identified up-regulated pathways that were previously suggested as leading to PTB like immunological and inflammatory paths. New pathways that were not identified in preterm like the hemopoietic path appeared up-regulated in preterm membranes. A group of 18 down-regulated genes discriminates between term and severe preterm cases. These genes potentially characterize a severe preterm transcriptome pattern and therefore are candidate genes for understanding the syndrome. Some of the down-regulated genes are involved in the nervous system, morphogenesis (*WNT-1, DLX5, PAPPA2*) and ion channel complexes (*KCNJ16, KCNB1*), making them good candidates as biomarkers of PTB.

The identification of this DE gene pattern may help to develop a multi-gene disease classifier. These markers were generated in an admixtured South American population where PTB has a high incidence. Since genetic background may impact differentially in different populations it is mandatory to include populations like South American and African ones that are usually excluded from high throughput approaches. These classifiers should be compared to those in other populations to get a global landscape of PTB.

## Introduction

Preterm birth (PTB), defined as the delivery of an infant before 37 weeks of completed gestation, is a worldwide health problem and remains the leading cause of global perinatal morbidity and mortality [1–3]. PTB is a complex, multifactorial syndrome comprised of multiple clinical subtypes that can be defined as either ‘spontaneous’ or ‘medically indicated’ [4,5]. 70 % of PTB cases are idiopathic; 5 % are due to the spontaneous onset of labor (sPTB) and the remaining 25 % are consequence of the preterm premature rupture of membranes (PPROM) [5,6]. PTB has also been stratified according to gestational age (GA); neonates born between 24 and 33 weeks (severe PTB) are at higher risk of death, and diseases later in life, than moderate PTB (GA between 34-36 weeks). It has been speculated that PTB from different GA groups has diverse causes and/or pathological mechanisms [6]. Regardless of PTB subtype, current therapies are not successful in prolonging time to birth once labor has been initiated [3].

PTB occurs as a result of the interaction of both, genetic and environmental components, and it constitutes a complex multifactorial syndrome [3,7]. Genetic architecture of pregnancy and PTB has proved challenging not only because of the multiple causes of PTB but also because of the numerous maternal and fetal gestational tissues that must interact to facilitate parturition [3,8]. These tissues include decidua, myometrium, cervix and maternal blood originating from the mother and villous placenta, fetal membranes (chorion and amnion), umbilical cord, and fetal blood originating from the fetus [5].

The main etiological factors related to PTB are inflammation, hemorrhage, activation of maternal or fetal hypothalamic-pituitary axis, immune dysregulation, distension of the myometrium and cervical insufficiency [1,3,7,9]. All these process have diverse and distinctive ways of initiating labor but may share a common pathway that end in the release of mediators that stimulate myometrial contraction, degradation of extracellular matrix components, inflammation and apoptosis. Consequently, these processes promote membrane rupture, cervical ripening, and uterine emptying resulting in PTB [1,9]. Studies of the transcriptome of those tissues can help to develop a molecular landscape of preterm labor and improve the understanding of the physiology and pathology of term and preterm parturition. Specifically, the study of the transcriptome of the membranes at the site of rupture in PTB may indicate shared genes of those pathways.

RNA sequencing (RNA-Seq) is a potent technology for the analysis of transcriptome that allows a comprehensive characterization of gene expression [10]. The published RNA-Seq studies on human labor have been restrained to normal term pregnancies and confined to placenta at different GA [11–13]. More placental gene expression data is available from experiments based on microarrays [14], but most of them are concentrated on preeclampsia [11].

As it was mentioned, GA determines diverse pathological mechanisms of PTB [6]. The genetic background seems to play a more relevant role in severe PTB neonates than in moderate PTB [15]. Modifications of chorioamniotic expression, that end in severe PTB should be more drastic than in term delivery or even moderate PTB. Therefore, we decided to focus on the transcriptome of severe PTB chorioamniotic tissues in an attempt to find DE genes with more biological significance. To validate the principal pathways found in this study, our results were compared with data reported in the previous literature. The ultimate goal of the study is to obtain an expression signature of PTB.

## Materials and Methods

### Patient recruitment

Controls and cases were term and preterm deliveries, respectively, from unrelated offspring of women in labor-receiving obstetrical care at the Pereira Rossell Hospital Center, Montevideo, Uruguay. Preterm tissues were collected from pregnancies complicated by spontaneous PTB before 33 weeks of gestational age (GA) while term chorioamnion tissues were obtained from no complicated pregnancies delivered after 37 weeks of GA. None of the patients presented multiple gestations, fetal anomalies, eclampsia, nor C-surgery delivery.

Maternal demographic characteristics were collected through questionnaires filled by mothers after delivery. Clinical and obstetric data were obtained from the Perinatal Information System which consists of basic perinatal clinical records developed by the Latin American Center for Perinatology (CLAP) from WHO/PAHO [16].

### Ethics statement

The School of Medicine’s Ethics Committee of the University of the Republic, (Comité de Etica de Facultad de Medicina-UDELAR) Uruguay approved the study protocol and an informed written consent was obtained from mothers prior to collection of biological material.

### Chorioamnion tissue collection, RNA extraction, and sequencing

Samples from 35 deliveries were collected within 30 minutes post-delivery. The amnion and chorion were obtained from the extraplacental membranes (reflected membranes), which provide a purer source of the fetal membranes. A portion of 1 cm^2^ of chorioamniotic membrane surrounding the exact place of membrane rupture for each subject was collected and immediately frozen in liquid nitrogen or submerged in RNA later solution (Qiagen). All samples were later stored at −80°C until laboratory procedures were performed. Samples were grinded into a fine powder in liquid nitrogen with a precooled pestle and mortar and subjected to RNA extraction.

Total RNA was extracted from each sample using a Trizol RNA extraction protocol that produces mRNA-enriched purification. Quality and concentration of RNA products were determined by UV-absorbance spectrophotometry (Nanodrop technologies Inc. Wilmington, DE, USA). The integrity of RNA molecules was checked using a 2100 Bioanalyzer platform (Agilent Technologies). We selected eight RNA samples (4 cases and 4 controls) for paired-end sequencing based on matching mother`s age, fetus sex and socioeconomic status. All held RNA Integrity Numbers (RINs) of 8 or higher. Total RNA samples were shipped to Macrogen Inc. (Seoul, South Korea) under recommended RNA submission conditions. On arrival samples were further assessed for RNA integrity on using a 2100 Bioanalyzer platform (Agilent Technologies). Messenger-RNA (mRNA) content was purified from total RNA with the PolyATract mRNA Isolation System II (Promega Inc.) and copied to cDNA molecules using Illumina TruSeq RNA Sample Preparation Kit v2. One such cDNA library was constructed for each specimen; libraries were subjected to massive sequencing following Illumina HiSeq2000 protocol (Illumina Inc.; www.illumina.com). A read was defined as a 100 bp cDNA fragment sequenced from both ends (paired-end). The data have been deposited in Sequence Read Archive (NCBI) and are accessible through SRA Series accession number SRP139931. A subset of extracted RNAs, making up a total of 15 terms and 9 severe PTB samples including the sequenced samples, were used to validate the results by real-time quantitative-PCR.

### Mapping reads to the reference genome

Obtained reads were trimmed and clipped for quality control in Trimmomatic v0.32 [17], and checked for quality using FastQC v0.11.2 [21]. Reads were then aligned to the GRCh37 reference genome using Tophat v2.1.0 [22] and Ensembl annotations [23], as derived from Ensembl Release 75.

Only for data visualization purposes, gene read counts were transformed by the regularized logarithm (*rlog*) [24]. This transformation removes the dependence of the variance on the mean and normalizes count data with respect to library size.

### Identification of differentially expressed (DE) genes and gene set enrichment analysis

HTSeq-count with parameters m = union, s = no, and t = exon was used to produce raw read counts of expression for each gene [25]. Differential expression analysis on the gene level was performed in R-packages DESeq2 v1.16.1[24], edgeR v3.18.0 [26] and Cuffdiff v2.2.1 [27]. P-values were adjusted for multiple testing using the Benjamini and Hochberg’s ("BH") approach [28] to control the false discovery rate (FDR) [23]. A gene was considered as expressed if it had more than 5 aligned reads. Genes were identified as DE with the following criteria: absolute log 2 fold change > 2 and FDR-adjusted p-value < 0.05 for multiple hypothesis testing (method=“BH”). With the aim of reducing false-positive hits, we required a gene to be selected by all three methods to be considered as DE.

To understand the underlying biological processes, functional annotations of DE genes were performed using GAGE and GOSeq packages of Bioconductor 3.5 [29,30]. GOSeq method considers the effect of selection bias in RNA-Seq data that can arise as a result of gene length differences [18]. All gene mapping was performed using the biomaRt R package [31]. We looked for enrichment for genetic association with KEGG pathways and Gene Ontology (GO) terms, and for GO terms that were supported by at least 3 analyzed genes. Multiple testing was adjusted using BH approach, and enrichment was declared if BH adjusted p-value was less than 0.05.

### Validation of RNA-Seq results by assessment of gene expression by real-time PCR (qRT-PCR)

Six genes were selected for the confirmation of DE gene data by qRT-PCR; Interleukin 1 beta (*IL1B*), lipocalin 2 (*LCN2*), macrophage receptor with collagenous structure (*MARCO*), caspase 5 (*CASP5*), serpin family A member 1 (*SERPINA1*), and TNF superfamily member 15 (*TNFSF15*). These gene’s expression values were evaluated on the 15 term birth and 9 severe PTB collected samples.

Total RNA of 1µg from each sample was used to synthesize the first-strand cDNA using Superscript II RT reagent kit (Invitrogen) and oligo(dT)_20_ primers. qRT-PCR amplifications were performed in a Corbett Real-time thermocycler (Qiagen), using Biotools SYBR Green kit (Biotools, Spain). The relative expression ratio of a target gene was calculated as described in the 2-DDCt method [32].

Reaction mixtures contained 10 µl of QUANTIMIX EASY kit (Biotools; #10606-4153), 0.5 µM of each forward and reverse primers, 1 µl of cDNA sample, and deionized water to a final volume of 10 µl. The following conditions were used: 95°C for 5 min and 40 cycles at 95°C for 15 seconds, followed by 1 minute at 63°C. After amplification a melting step was performed, with a rise in the temperature from 72 to 90°C with continuous acquisition of fluorescence.

A positive and negative (non-template control, NTC) control was added to each PCR reaction. Each sample was assessed in duplicate and the %CV between the duplicates was < 2%. All primers sequences for the validated genes were designed to span exon-exon junctions to minimize the potential of amplifying genomic DNA (S1 Table). Amplification efficiencies of primers were within a range of 90% and 110% efficiency, and primer specificity was assessed by the presence of a single temperature dissociation peak. The glyceraldehyde-3-phosphate dehydrogenase (*GAPDH*) gene and the TATA-box protein (*TBP*) gene were chosen as the reference genes to estimate the relative quantification. The arithmetic mean of GAPDH and TBP genes was used as the reference. The relative gene expression was calculated using the method 2^−ΔΔCt^ with the term group as the control group [32].

### Validation of RNA seq results by comparison with previous preterm transcriptomic studies

In order to determine if the results presented here reflect the PTB mechanisms that are previously reported in the literature, a Pubmed search was performed using these search terms: [preterm AND (transcriptome OR transcriptomic)]. The electronic search was performed on March 19^th^, 2018 in PubMed with no restrictions, to identify all articles relating to DE genes in all gestational tissues. The results were analyzed based on 5 inclusion criteria: published in English, original research, human chorioamnion tissue samples, DE between term and PTB and candidate gene list assembled. All articles were cross-checked with database search results to find any additional available transcriptome data.

### Statistical analysis and data visualization

Maternal demographic and clinical characteristics of term and severe preterm study groups were compared using the Student’s t-test for between-group comparisons of continuous data. For comparisons of categorical data, the Chi-square test was used. All statistical analyses were performed using the R environment for statistical computing version 3.4.3 [19]. After multiple test correction, a p-value <0.05 was used to consider statistical significance. The ggplot2 R package was used to create plots [19].

## Results

Demographic and clinical characteristics of the term and severe preterm groups are presented in Table 1. There were significant differences between cases and controls for the gestational age and prenatal medical appointments. The remaining assessed variables did not present significant differences between groups.

**Table 1.**
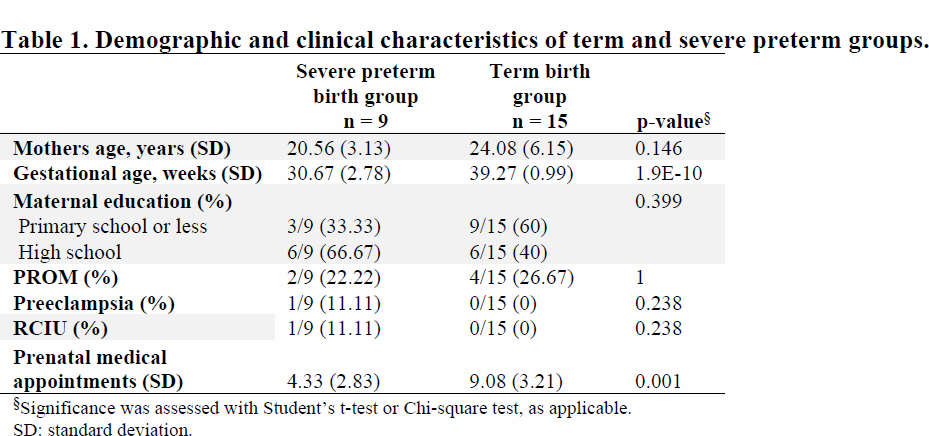
Demographic and clinical characteristics of term and severe preterm groups.

### Exploration and evaluation of overall gene expression

Genome-wide gene expression in chorioamnion tissue samples was evaluated using RNA-Seq technology. A summary of the sequencing read alignments to the reference genome is presented in S2 Table. Illumina sequencing was effective at producing large numbers of high quality reads from all samples. On average, 93.7% of the total reads were mapped successfully. Among the aligned reads, 91.5% were mapped to unique genomic regions.

A Principal Components Analysis (PCA) was performed on the gene expression rlog transformed values (Fig 1A). A high portion of the overall variance was accounted for by the first two principal components of this model (PC1, 64 %; PC2, 22 %). The PCA shows term samples as a clearly packed group, while severe preterm samples showed greater variability.

**Fig 1.**
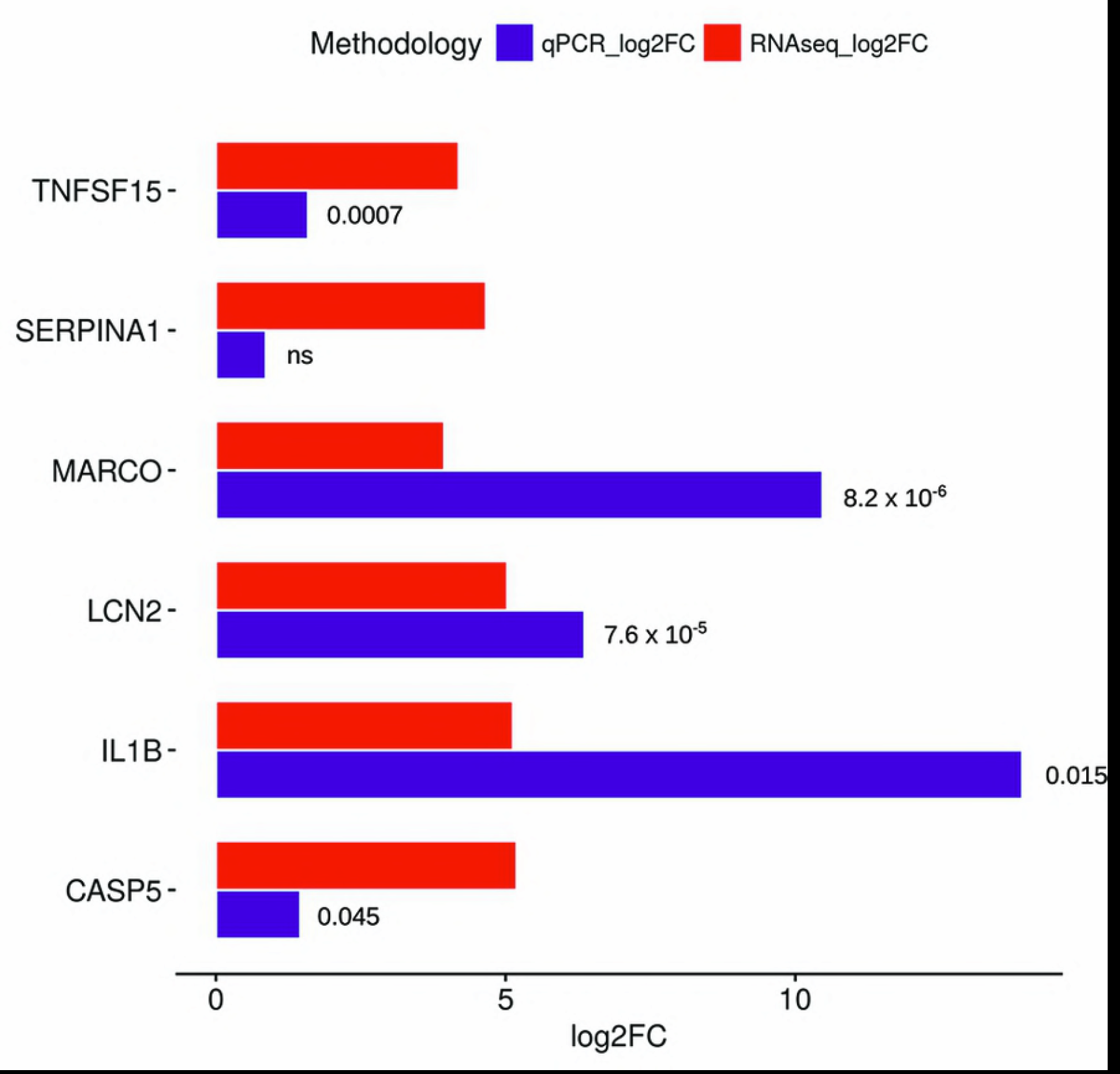
Overall gene expression of fetal membrane transcriptomes. (A) Principal Component Analysis (PCA) plot. (B) Heatmap of regularized log (rlog) normalized values of expression levels and hierarchical clustering for the 270 DE genes (FDR ≤ 0.05 and absolute logarithm fold change ≥ 2). The condition of each sample is denoted at the top: red for term samples and blue for severe preterm samples. Rlog values are coded on the blue-to-orange scale (high expression: orange; low expression: blue). Blue dots on the gene’s dendogram represent 3 major groups according to gene expression patterns.

A total of 270 genes were DE simultaneously by the 3 methods employed (FDR ≤ 0.05 and absolute logarithm fold change ≥ 2) (S1 Fig). Within the DE genes, 252 were up-regulated and 18 were down-regulated in the severe preterm compared to the term births. The full list of DE genes detected (FDR ≤ 0.01 and fold change ≥ 2) can be found in S1 Supplemental File.

When visualizing the *rlog* transformed read counts of the 270 significantly DE genes, control samples have a homogenous expression pattern (Fig 1B). Notably, several of the DE genes were related to immune-related functions such as interleukin 24, CXC motif chemokine ligands and Tumor Necrosis Factors signaling pathways. A hierarchical clustering analysis clusters genes in 3 groups. Remarkably, one of the clusters is a group of 18 genes which has a notably discordant expression pattern for term and severe preterm samples. This group is constituted by the aforementioned 18 down-regulated genes (Fig 2), which potentially characterize a severe preterm transcriptome pattern, and therefore are candidate genes for the understanding of the syndrome. These genes include potassium voltage-gated channel subfamily members, genes involved cell adhesion functions, the placental alkaline phosphatase and a Wnt family member. This gene cluster is related to GO terms involved in the nervous system, organ morphogenesis, and ion channel complexes, but no term remained as enriched with statistical significance in the group after BH p-value correction.

**Fig 2.**
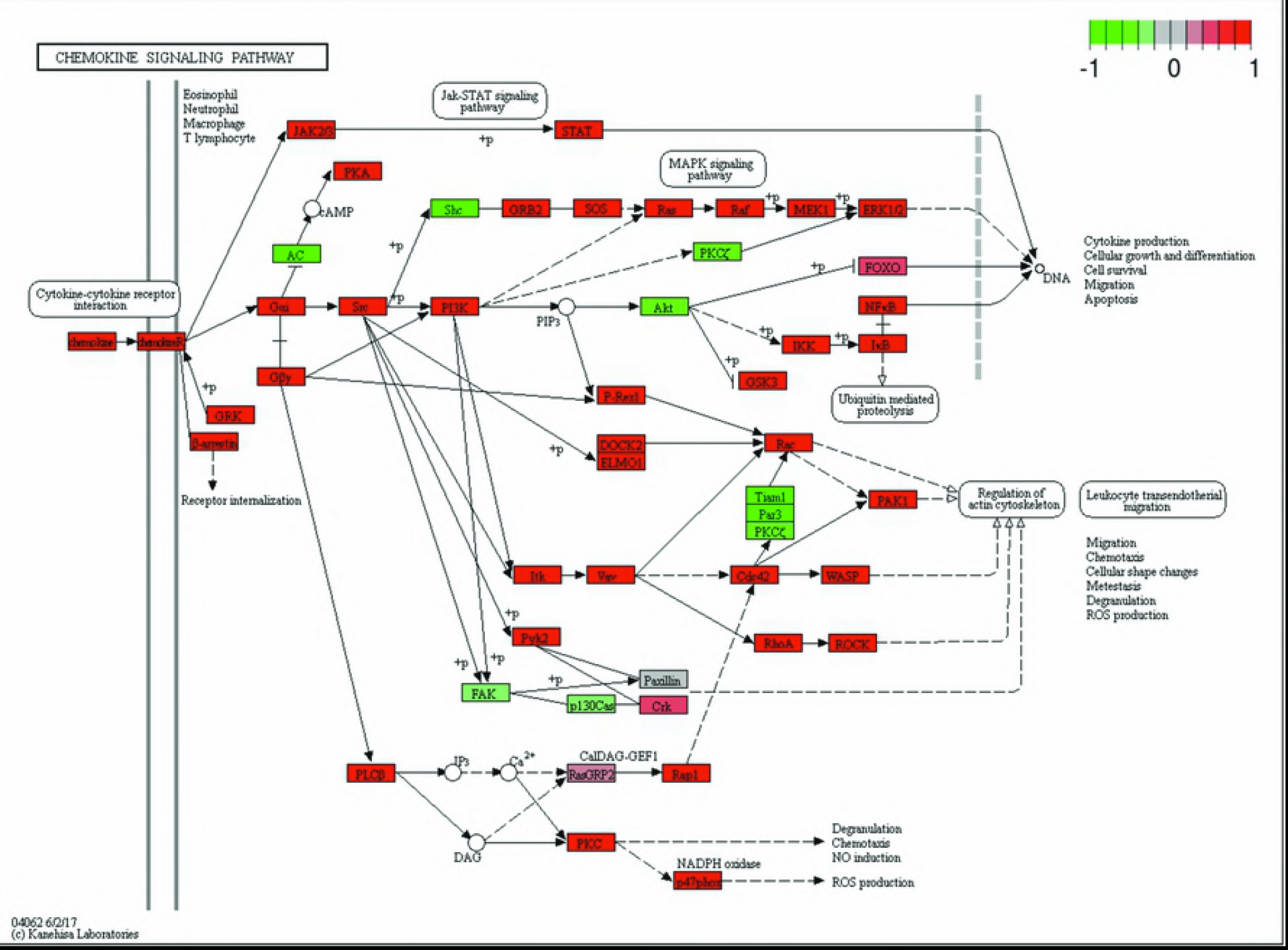
Gene expression levels of the 18 down-regulated genes. Heatmap of regularized log (rlog) normalized values of expression levels and hierarchical clustering for the 18 down-regulated DE genes. The condition of each sample is denoted at the top: red for term samples and blue for severe preterm samples. Gene symbols are stated at the right. *rlog* values are coded on the red-to-green scale (high expression: red; low expression: green).

The remainder two groups are constituted by 147 and 105 genes. Mainly, they have genes related to chemokines, TNF family members, coagulation factors, transmembrane proteins. A GO term enrichment search revealed 508 and 301 terms, respectively, were statistically significant (BH corrected p-value < 0.05). These terms were mostly immune and inflammatory response and immune system related terms.

### Gene Set Enrichment Analysis using Gene Ontology

In an effort to identify processes and pathways that could be regulated differentially between severe preterm and term groups, we performed a Gene Set Enrichment Analysis (GSEA), as implemented in goseq R package. Genes that showed a FDR ≤ 0.05 were tested against the background set of all expressed genes with ENSEMBL annotations. In this sense, for the enrichment analysis comparing the gene expression between severe preterm and term, 270 significant genes were tested against 18,675 background genes. The significant enrichment of Gene Ontology (GO) terms was tested using the Wallenius approximation and applying multiple hypothesis testing correction (BH). Only terms with 3 or more statistically significant genes were considered.

A total of 1886 GO terms were found enriched among the DE genes (FDR < 0.05), corresponding to Biological Processes (1495 terms), Cellular Component (195) and Molecular Function (196) (Fig 3A). Within Biological Processes category, *immune response*, *immune system process* and *defense response* terms were the most significantly enriched with DE genes. Likewise, the Cellular Component category subdivided annotated sequences into plasma membrane, cell periphery, and *cytoplasmic vesicle part* as the most represented terms. Within the category Molecular Function, the three principal groups were: *signaling receptor activity*, *rec*eptor activity, and transducer activity.

**Fig 3.**
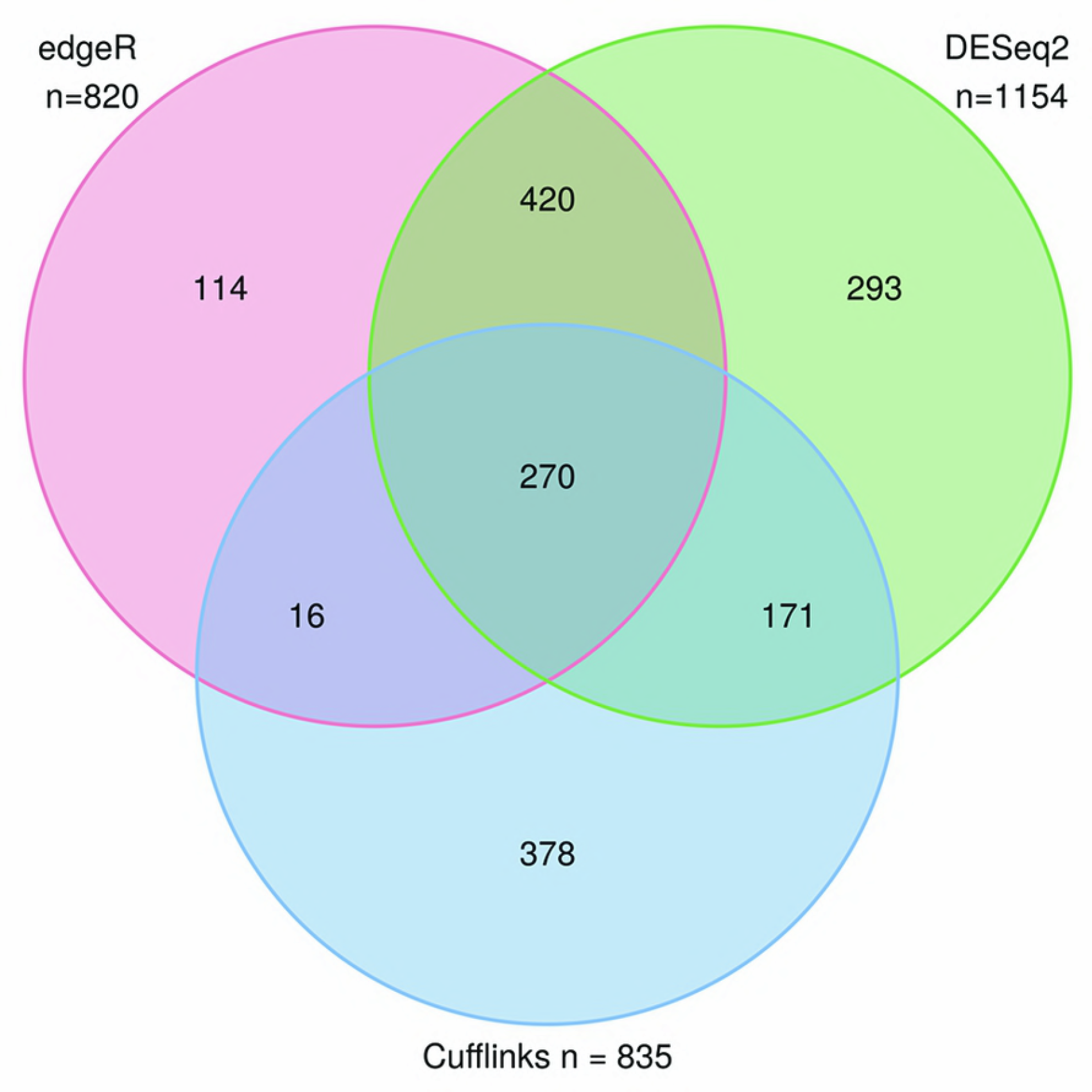
Enriched GO categories (A) and KEGG pathways (B) of genes in severe preterm birth. Enrichment analysis of Biological Process GO categories (A) and KEGG pathways (B) of DE genes in severe preterm/term comparison. Top 10 most significant terms or pathways are shown. The x-axis represents the enrichment value, as the logarithm of adjusted p-value (FDR). The number of genes identified in each pathway is reported inside brackets. All represented GO terms are significantly enriched, with an adjusted p-value <0.01.

When analyzing KEGG pathways, the most significant up-regulated pathways are Chemokine signaling pathway and Hematopoietic cell lineage (both with FDR = 7.038 x 10^−09^; Fig 4). Several relevant processes for PTB were up-regulated such as those involved in intracellular signaling: Toll-like, NOD-like, T cell and Jak-STAT signaling pathway (Fig 3B). No statistically significant down-regulated pathways were found.

**Fig 4.**
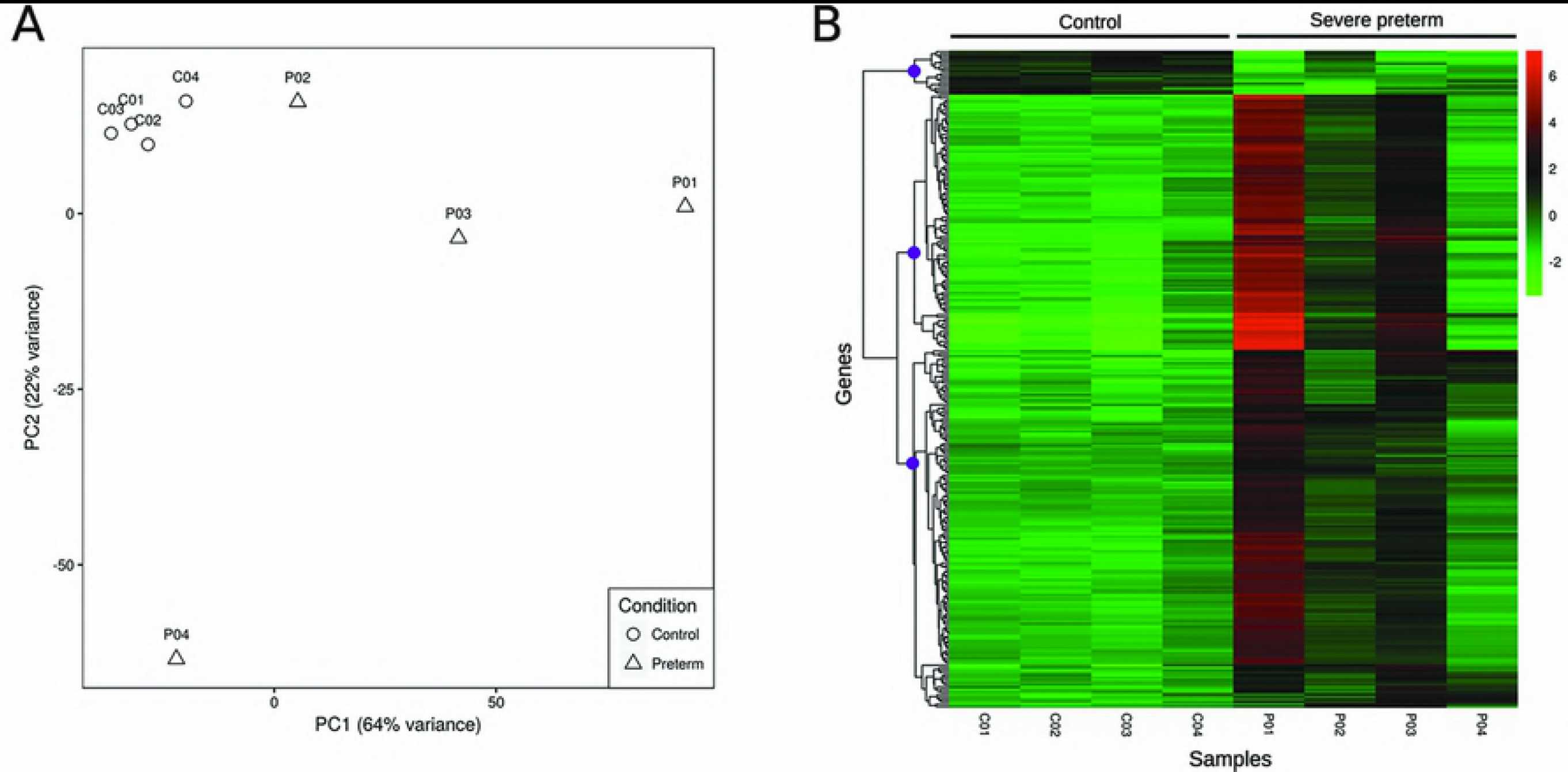
KEGG diagram of the chemokine signaling pathway. Genes overexpressed in preterm birth fetal membranes are red, and down-regulated genes are green.

### Validation of gene expression by qRT-PCR

To validate genes found to be significant in the RNA-Seq analysis, six DE genes were selected: *IL1B, LCN2, MARCO, CASP5, SERPINA1* and *TNFSF15*, and their expression was assessed by qPCR. The RNA-Seq analysis showed all genes were significantly up-regulated in PTB. Fig 5 displays the logarithm fold differences in gene expression measured by both RNA-Seq and qRTPCR. The six genes displayed similar patterns of mRNA abundance with both methods. Furthermore, *LCN2* and *MARCO* had statistical significance differential expression between severe preterm and controls when analyzing qPCR expression data.

**Figure 5.**
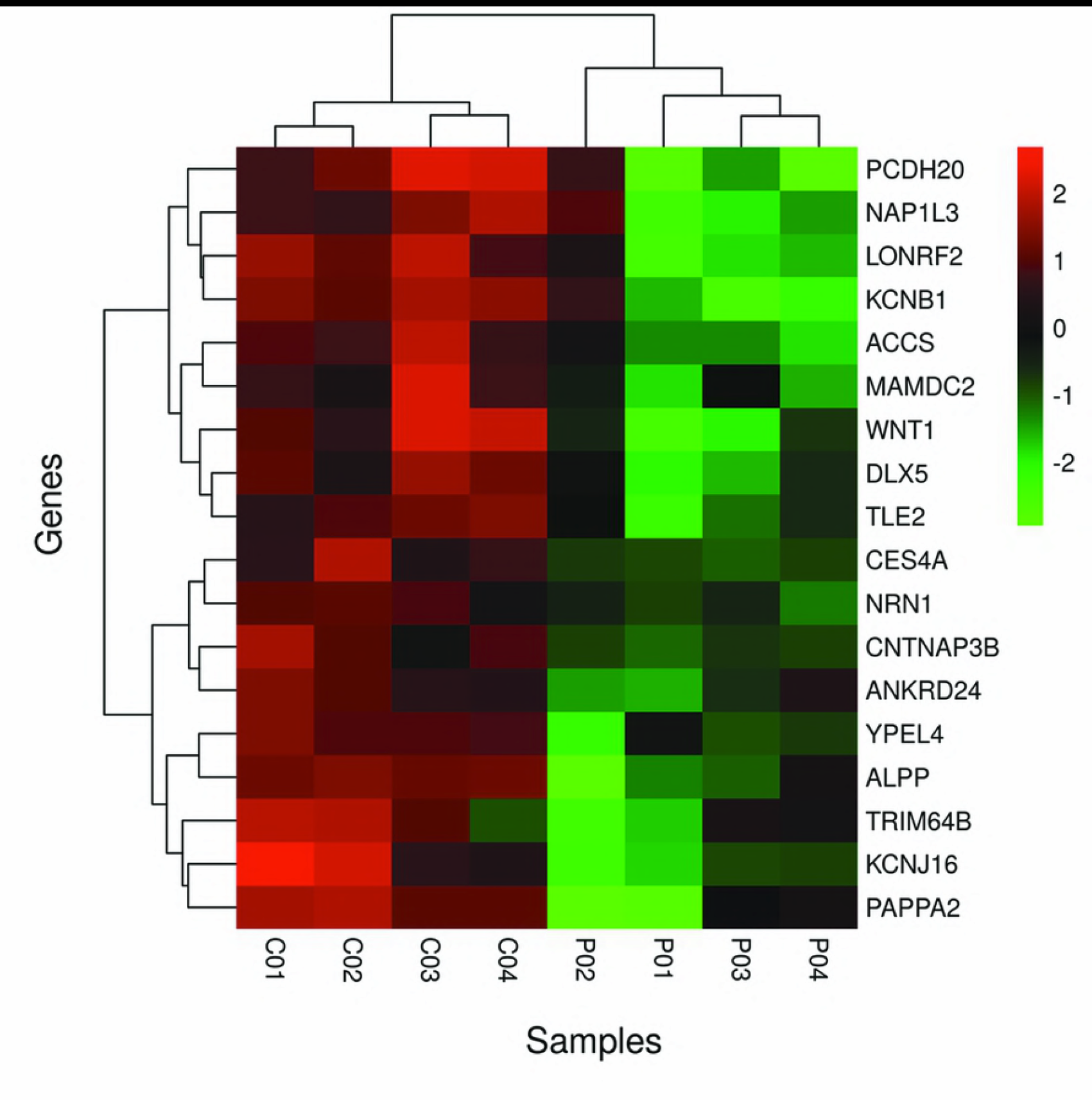
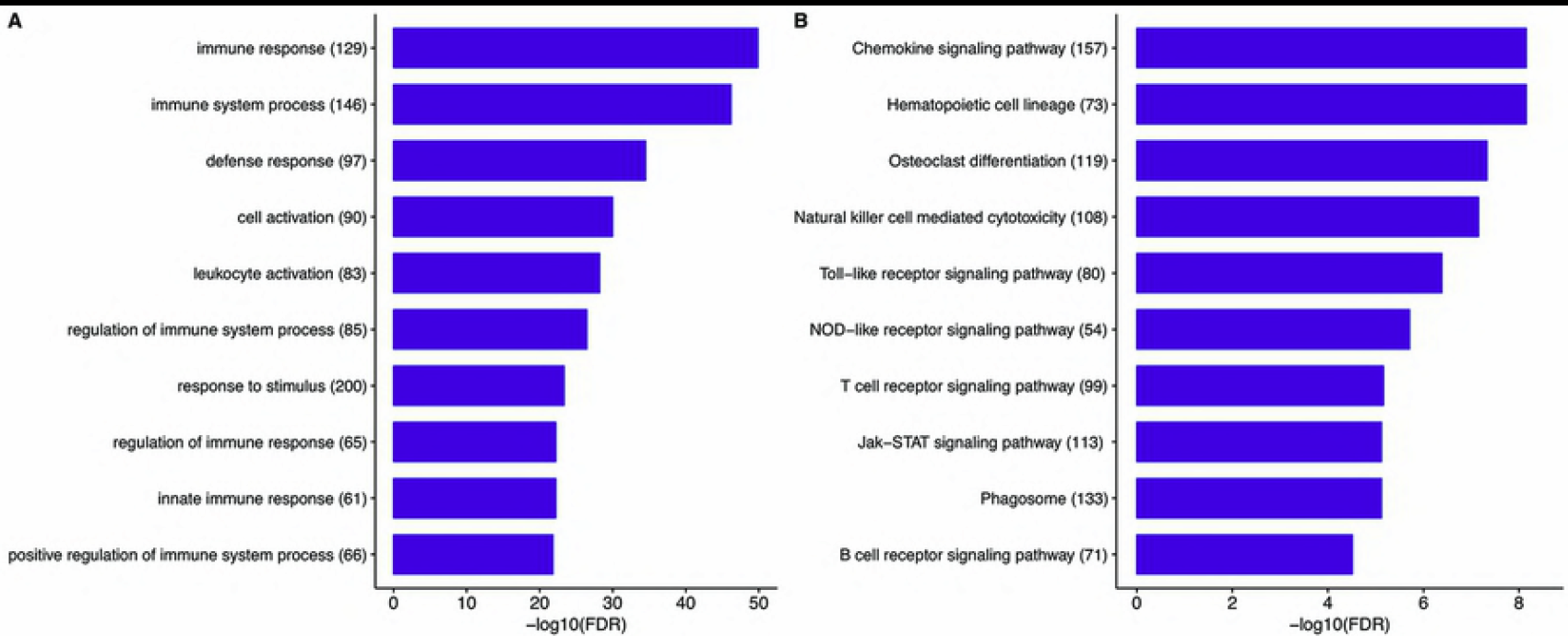
Validation of overall gene expression. Fold changes of five DE genes measured by RNA-Seq (red) versus qRT-PCR (blue). Significance values for qRT-PCR fold-changes are presented. IL1B: Interleukin 1 beta, LCN2: lipocalin 2, MARCO: macrophage receptor with collagenous structure, CASP5: caspase 5, SERPINA1: serpin family A member 1, and TNFSF15: TNF superfamily member 15. ns: non-significant (adjusted p value>0.05).

### Validation of RNA seq results by comparison with previous preterm transcriptomic studies

The results present in this study were compared with those previously reported in the literature. A PubMed search using the terms [preterm AND (transcriptome OR transcriptomic)] yielded 101 results. We collected the 101 abstracts and after analysing them, we found that most studies focused on preeclampsia, placental villi tissue, and animal models or maternal or cord blood. Six studies used RNAseq; two focused in cord blood [20,21], two on animal models [22,23] and two on the transcriptome of normal labor [12,24].

Only one study met all inclusion criteria mentioned above and included the transcriptome of chorion and amnion membranes and was included to match the results obtained in this work [25]. In this study, they used the microarray technology and found 50 genes that identified PT labor phenotype, seven of which we found as DE in this study (Table 1). The gene’s change direction of gene expression matches between studies in all cases [25].

**Table 2.** DE genes simultaneously found in Bukowski et al. (2017) and this study.

| Gene Symbol | Gene | HGNC ID | Change direction |
| --- | --- | --- | --- |
| BCL2A1 | BCL2 related protein A1 | HGNC:991 | up-regulated in PTB |
| CCRL2 | C-C motif chemokine receptor-like 2 | HGNC:1612 | up-regulated in PTB |
| CHI3L1 | chitinase 3 like 1 | HGNC:1932 | up-regulated in PTB |
| CXCL2 | C-X-C motif chemokine ligand 2 | HGNC:4603 | up-regulated in PTB |
| GPR84 | G protein-coupled receptor 84 | HGNC:4535 | up-regulated in PTB |
| SAMSN1 | SAM domain, SH3 domain and nuclear localization signals 1 | HGNC:10528 | up-regulated in PTB |
| SLC16A10 | solute carrier family 16 member 10 | HGNC:17027 | up-regulated in PTB |

## Discussion

Although PTB is responsible for most neonatal deaths and morbidity [26], there has been a shortage of studies that permit to create a distinctive expression footprint of the pathology and consequently achieve improvements in its prevention, diagnosis, and treatment. To overcome this gap in knowledge, we analyzed the gene expression of the chorioamnion membranes from severe PTB and term newborns by RNA-seq.

In this study, we present a distinct transcriptome pattern of severe PTB. Our study revealed 270 DE genes between both conditions. A number of this magnitude may partially explain the lack of previous molecular signatures for PTB as well as the difficulty to identify biomarkers and potential therapeutic interventions [4,39]. Several pathways and genes are involved in labor and most of them would be either turn on or off during pathological conditions [4,16,37,40].

It has been proposed than pregnancy is maintained by down-regulation of chemokines at the maternal-fetal interface [25]. Removal of the down-regulation results in term birth while in the case of PTB, the activation of multiple pathways of the immune system overrides this down-regulation [3,22,25,27–30]. In concordance with this idea, we found that up-regulated genes were mostly related to immune response (e.g. interleukin 24, CXC motif chemokine ligands or Tumor Necrosis Factors). Moreover, we found that severe preterm labor has a significant enrichment of immune processes in comparison to term labor. Gene ontology and KEGG pathway enrichment analyses indicated that inflammatory and immune paths were enriched in severe preterm samples (Fig 3). These findings are in agreement with the mentioned studies, demonstrating that preterm labor is associated with an extensive immune response activation in several gestational tissues. In fact, the incidence of PTB in association with inflammation is higher earlier in pregnancy [31], so the overexpression of inflammatory and immune pathways in severe PTB is consistent with the results reported in the literature.

Interestingly, two of the overexpressed genes found in this study were previously proposed as biomarkers of PTB because their concentration in amniotic fluid was higher in PTB when compared to term labor: C-X-C motif chemokine ligand 10 (*CXC10*) gene and matrix metallopeptidase 8 (*MMP-8*) [32–35]. Consequently, these biomarkers can be analyzed in amniotic fluid and provide information to prevent PTB. CXCL10 is related to maternal anti-fetal rejection, in the same way as it is with allograft rejection [34]. In this context, *CLXC10* overexpression in PTBs may be leading to premature labor by triggering a fetal inflammatory response. MMP‐8 is known to decrease the integrity of the cervical and fetal membrane extracellular matrix [36]. MMP-8 may alter the membranes extracellular matrix and permit the passage of bacteria into the endometrium, leading to bacteria-induced PTB [37].

A small group of 18 genes clustered together and discriminates between term and severe preterm cases (Figure 2B). All of them are down-regulated. These genes potentially characterize a severe preterm transcriptome pattern and therefore are candidate genes for understanding the syndrome. None of these down-regulated genes were previously proposed in the literature as preterm birth biomarkers. However, placental alkaline phosphatase (*ALPP*) and pappalysin 1 (*PAPPA*) gene are included in a group of nine placental genes that together could predict GA in a recent pilot study [38]. ALPP and Pappalysin 2 (*PAPPA2*), a closely related gene to PAPPA, are among the 18 down-regulated gene cluster found here. Also, PAPPA2, a type of metalloproteinase, with a typical high gene expression in the placenta [39], was found DE in placenta tissue between spontaneous PTB and term birth [40].

The low number of down-regulated genes made difficult to establish statistically significant pathways: only two from the 18 clustered genes are included in the same KEGG pathway. WNT1 and *DLX5* genes belong to the pathway: “Signaling pathways regulating pluripotency of stem cells” (hsa04550). In term births, this pathway increases its expression at the end of pregnancy [41]. *WNT1* gene belongs to the WNT signaling pathway, which is involved in cell proliferation, development, and tumor progression. WNT signaling pathway inhibition could contribute to miscarriages and PTB, via decreasing trophoblast proliferation and invasion [42]. Notoriously, *WNT2* gene, which is typically overexpressed in placenta, was not DE in our data. *DLX5* is a member of a homeobox transcription factor gene family. Homeobox genes regulate embryonic as well as placental development [43]. They are widely expressed in the human placenta. A potential role of DLX5 may be related to neurobehavioural development and to regulating stem cell function, both important for normal placental development [43]. Also, *DLX5* is up-regulated in preeclamptic placentas, presumably due to an altered methylation at the *DLX5* locus in preeclampsia that results in loss of imprinting [44]. A decrease in expression later in gestation was found for this gene in human placenta, as a response to an increase in gene body methylation over gestational age [43]. Our results indicate that *WNT1* and *DLX5* expression are lower in fetal membranes of severe PTBs than in term membranes. In this context, we speculate that the down-regulated genes may act as maturation genes setting the gestation timing.

These putative maturation genes may be very difficult to discriminate from those leading to preterm birth. A major issue in the transcriptomic study of PTB in humans is the inability to collect healthy control tissue at the same gestational age (GA) to compare with pathologic preterm tissue. Consequently, DE genes between PTB and term reflect differences of both, GA and the pathological event of preterm labor. We are aware that as a consequence of the design of our study PTB and GA responsible genes are overlapped. By comparing human sPTB and term transcriptomes with GA-matched control transcriptomes from a closely related species (macaques), Eidem *et al. [45]*, identified a low number of distinctive promising sPTB-specific candidate genes or potentially related to GA effects. They found similar functions of PTB and GA genes so they speculated that the effects of sPTB and GA do not correspond to biologically distinct processes [45].

Whether PTB is a labor that occurs too soon or it represents a pathological event with a new group of expressed genes is still an open question. The present transcriptome of chorioamnion membranes does not resolve the controversy but it adds information that would help to complete the PTB expression scenario. On one hand, our results confirmed that immune response pathways are up-regulated in PTB. On the other hand, this RNA-seq approach allowed to identify up-regulated genes and pathways not previously reported as associated with PTB (hematopoietic cell lineage pathway, osteoclast differentiation). Also, we found that some of the down-regulated genes are involved in the nervous system, morphogenesis (WNT-1, DLX5, PAPPA2) and ion channel complexes (KCNJ16, KCNB1). It is difficult to speculate on the importance of those genes and pathways in PTB; for example ion channels are involved in a wide spectrum of different functions that eventually conduct to labor or PTB including implantation [46], Ca^++^ signaling [47], smooth muscle contraction [48] and inflammation [49]. Regarding the possibility that hematopoietic cell lineage pathway is connected with PTB, it is known that the placenta produces hematopoietic stem cells (HSCs) [50,51]. The placenta can generate HPCs and cause their expansion but not differentiation. Placental HSC pool was first considered to appear early in gestation and then to decrease while the development of the HSC reservoir in the liver [51,52]. Subsequent studies demonstrated that mid-gestation placenta also harbors a large pool of pluripotent hematopoietic stem cells that have the capacity to self-renewal [50][53]. Moreover, hematopoietic progenitors were also found in full-term placentas in a low percentage of lineage-committed cells [54]. Since the hematopoietic monocyte-macrophage lineage produces the osteoclasts, hematopoietic cell lineage and osteoclast differentiation pathways are connected. Although *a priori* both pathways look far away from term labor or PTB, they could represent interesting unexplored paths to preterm. Nevertheless, all of the mentioned genes and biological process open new lines of research making them good candidates as biomarkers of PTB.

A literature search to validate these genes revealed that, surprisingly, there are not many high-throughput studies analyzing chorioamniotic tissues and none with RNA-seq technology. To our knowledge, this is the first RNA-seq study of chorioamniotic tissue in preterm and term birth. When comparing results from Bukowski et al. [25] only 7 genes are shared with our study. All these genes may be in one way or another related to PTB: *CCRL2* and *CXCL2* belong to inflammatory pathways [55,56], GPR84 is a receptor of fatty acids also related to inflammation [57]. BCL2A1 and CHI3L1 were recently associated with chorioamnionitis in monkey and human [58], SLC16A10 is up-regulated in the mid-gestational placenta [59] and *SAMSN1* is a poorly characterized gene that may be involved in differentiation of lymphocytes B [60]. The fact that these subsets of genes overlap in both studies using different experimental techniques stresses that they should be a target of further research of PTB.

One interesting finding is the homogeneity for gene expression in the control samples. After a linear reduction of the variability for more than 30000 expressed genes, the control samples cluster together in a small group. The dispersion of the samples in the PCA analysis is lower compared to other tissue expression profiles using the same exploratory analysis [61]. In contrast, PTB chorioamniotic expression analysis revealed a more diverse group, further stressing the heterogeneity of this condition. This may be a consequence of the complexity of the syndrome characterization as it has been previously reported in the expression of other tissues related to PTB like the myometrium [62]. In concordance, Ngo et al. [38] developed a gene expression model based on GA that predicted time until delivery for full-term but that failed for preterm deliveries. This suggests that to predict PTB the classifiers must incorporate the several outlier physiological events that may lead to PTB [38]. Although we hypothesized that chorioamnion’s expression reflects a common pathway to most of the paths that end in PTB, it is possible that in different individuals diverse pathways participate making more difficult to define a landscape of PTB than one of term labor.

Finally, this study has been performed in an admixed population where PTB is frequent (approx. 10 %) indicating that it is a serious health public problem. Population-specific studies are relevant, as genetic background may impact differently in different populations, e.g. it is known that African American ancestry increases the risk of PTB [63–65]. Generally, there is a gap of information regarding specific populations except those coming from North America and Europe. With our literature search, we recovered only three out of 101 transcriptomic analyses based on populations from Latin American countries [34,66,67], stressing the fact that most studies do not include susceptible populations from this part of the continent.

All of the Latin American studies were performed with microarray analysis, meaning that the data shown in this work will be the first RNA-seq data available to contrast in future studies. Specifically, the Uruguayan population is admixed, with a predominance of European contribution, and a Native American and African contribution of 10.4% and 5.6%, respectively [68,69]. Common genetic variation affects expression variability [70], as well as the admixtured genome structure [71,72], stressing the necessity of population-specific transcriptome analyses or alternatively including diversity in high throughput population analyses.

## Conclusions

The use of state-of-the-art analytics by RNA-seq identified already reported genes and pathways as well as novel candidates involved in labor and PTB. The combination of up- and down-regulated genes permitted to develop a multi-gene disease classifier. These markers were generated in an admixtured population where PTB has a high incidence. Further studies including the testing of these classifiers using a second and larger independent sample are needed to validate these results. Moreover, these markers should be compared to other population studies to get a global landscape of severe PTB.

## Acknowledgements

We acknowledge Drs. Ariel Díaz, Natalia Cabrera and Pamela Grimaldi (Pereira Rossell Hospital Center) for their invaluable support in patient and control subject recruitment.

## Author Contributions

SP, BB and RS conceived and designed the experiments. CS assisted in sample collection. SP performed the experiments and analyzed the data. All authors discussed results. SP, BB and RS wrote the manuscript.

## Supporting information captions

**S1 Fig 1. Differentially expressed genes.** Venn diagram showing the number of differentially expressed genes identified by each of the three methods employed, when considering a FDR < 0.05.

**S1 Table. Primers used for quantitative PCR analysis**. Sequence of the primers used for qRTPCR

**S2 Table. Alignment statistics obtained by tophat2.**

**S1 File 1. Full list of differentially expressed genes**. (FDR ≤ 0.05 and fold change ≥éh8éõütüZtüõé in the severe preterm vs. term pairwise comparison.

